# Collagen hydrogel confinement of amyloid-*β* accelerates aggregation and reduces cytotoxic effects

**DOI:** 10.1101/711622

**Authors:** Laura W. Simpson, Gregory L. Szeto, Hacene Boukari, Theresa A. Good, Jennie B. Leach

## Abstract

Alzheimer’s disease (AD) is the most common form of dementia and is associated with the accumulation of amyloid-β (Aβ), a peptide whose aggregation has been associated with neurotoxicity. Drugs targeting Aβ have shown great promise in 2D *in vitro* models and mouse models, yet preclinical and clinical trials for AD have been highly disappointing. We propose that current *in vitro* culture systems for discovering and developing AD drugs have significant limitations; specifically, that Aβ aggregation is vastly different in these 2D cultures carried out on flat plastic or glass substrates vs. in a 3D environment, such as brain tissue, where Aβ confinement very likely alters aggregation kinetics and thermodynamics. In this work, we identified attenuation of Aβ cytotoxicity in 3D hydrogel culture compared to 2D cell culture. We investigated Aβ structure and aggregation in solution vs. hydrogel using Transmission Electron Microscopy (TEM), Fluorescence Correlation Spectroscopy (FCS), and Thioflavin T (ThT) assays. Our results reveal that the equilibrium is shifted to stable β-sheet aggregates in hydrogels and away from the relatively unstable/unstructured presumed toxic oligomeric Aβ species in solution. Volume exclusion imparted by hydrogel confinement stabilizes unfolded, presumably toxic species, promoting stable extended β-sheet fibrils. These results, taken together with the many recent reports that 3D hydrogel cell cultures enable cell morphologies and epigenetic changes that are more similar to cells *in vivo* compared to 2D cultures, strongly suggest that AD drugs should be tested in 3D culture systems as a step along the development pathway towards new, more effective therapeutics.

## 1. Introduction

Amyloids are defined as abnormal fibrous, extracellular, protein aggregates that accumulate in organs throughout the body. Native proteins misfold, and intrinsically-disordered proteins fold predominantly into a β-sheet conformation. Under certain conditions the β-sheet structure extends, leading to the formation of amyloid fibrils; this aggregation process either leads to loss of function of the native protein or gain of toxic function of the aggregated amyloid protein. There are currently 37 human amyloid proteins identified that are linked to neurodegenerative diseases and amyloidosis diseases [1, 2]. Alzheimer’s disease (AD) is the most common form of dementia [3] and is associated with the accumulation of amyloid-β (Aβ), and intrinsically disordered peptide between 39 to 45 amino acids long in its native form, whose aggregation has been associated with neurotoxicity in AD [4].

Since the first studies demonstrating a genetic link between Aβ and early-onset AD [5], investigators have targeted Aβ as a potential therapeutic strategy. Neurology has the third highest number of medications in drug development, yet there are only four unique drugs approved by the FDA for the treatment of AD [6]. Between 2002 and 2012 there were 413 clinical trials for AD. During that time only one drug was licensed by the FDA, a 99.6% failure rate [7]. Many AD drugs are Aβ targeting antibodies that demonstrated promise in pre-clinical trials and mouse models yet failed to show efficacy in human patients [8, 9].

There are a number of possible causes for the failure of Aβ targeting drugs in clinical trials. Aβ may not be causative in AD, but instead may simply be a side-product of the disease progression. The clinical trial design may be faulty in that drug intervention is administered in late-stage AD while Aβ toxic effects occur significantly before (potentially decades before) the onset of symptoms. Clinical endpoints are subjective, making it difficult to determine the efficacy of any intervention. Alternatively, the 2D tissue culture models used to screen drugs for their ability to attenuate Aβ toxicity are poor models for the *in vivo* 3D environment. Some or all of the above hypotheses may be valid; however, in this study, we set out to test the last hypothesis, that 2D tissue culture models are a poor model for Aβ toxicity testing.

*In vitro* models typically culture cells on a 2D surface to study the cellular response to Aβ [10–15]. However, 3D biomaterial scaffolds promote stem cell differentiation, cellular binding, and cell signaling that better represent *in vivo* cell behaviors compared to 2D cultures [16–24]. There is a growing trend across all of biotechnology and biopharmaceutical discovery teams to use 3D models of tissue as opposed to 2D tissue culture models, with remarkably good results [25–27]. 3D AD models have begun to be used to model the human cellular aspects that mouse models inherently cannot. [28–31]. Some of these 3D models have used iPSCs taken from an AD patient and differentiated them into neurons that produced Aβ and eventually tau tangles [30–32]. However, these models have yet to be used routinely for drug testing.

3D environments such as collagen hydrogels used in tissue culture models form a confined space that excludes solvent volume from soluble macromolecules [33–35]. Volume exclusion reduces the space available to an unfolded protein, thereby shifting the equilibrium towards a more compact protein conformation. For most proteins, the native state is a compact conformation [36–42]. However, with intrinsically disordered proteins and other aggregation-prone proteins, confinement promotes rapid aggregation [43–50]. Even for free, uncrosslinked crowding materials, the solvent exclusion effect has been observed experimentally [45, 47] and predicted from simulation [51, 52]. However, whether this effect leads to amorphous (off-pathway) aggregation [53, 54], or stabilization of on-pathway intermediates [55], is unclear as conflicting results have been reported [56]. Further, for hydrogels used as cell scaffolds (e.g., collagen) the influence of 3D environments on protein structure and aggregation is equally unclear.

From studies carried out in solution and 2D culture models, we know that Aβ aggregation is intimately tied to its cytotoxicity. Unstructured Aβ monomers associate with each other, forming unstable oligomers that are associated with neuronal toxicity [57–61]. Fibrils and other protofibril structures are thought to be less toxic [61–63]. While it is commonly held that the unstable toxic oligomers are an “on pathway” intermediate, there are some dissenting opinions [64]. Thus, a priori, it is difficult to predict the effect of 3D confined environments on the relative distribution of Aβ structures associated with aggregation and toxicity.

In this work, we tested the hypothesis that Aβ toxicity is altered for cells cultured in 3D hydrogels vs. 2D cell culture plates. We show that Aβ aggregates more readily in the 3D confined environment of a hydrogel. We suggest the differences in toxicity between 2D and 3D cultures are a result in a shift in chemical equilibrium from oligomer to fibril, and from toxic to less toxic species, in these environments. Finally, we propose that some clinical trials of AD drugs fail as a result of the differences in the biophysical environment in 2D culture and the *in vivo* environment; therefore, we suggest that future Aβ toxicity studies be conducted in 3D hydrogel cultures so that the biophysics of Aβ aggregation better approximates the *in vivo* scenario.

## 2. Materials and Methods

### 2.1. Beta-Amyloid Preparation

Human beta-amyloid (1-42) (Aβ) and scrambled Aβ (1-42) (Scr Aβ) (*AIAEGDSHVLKEGAYMEIFDVQGHVFGGKIFRVVDLGSHNVA*) was purchased from AnaSpec (Fremont, CA) and Genscript (Piscataway, NJ). HiLyte 488-labeled Aβ (1-42) (HiLyte Aβ) and FAM-labeled scrambled Aβ (1-42) (FAM Scr Aβ) were purchased from AnaSpec (Fremont, CA). All other unspecified reagents were purchased from Sigma Aldrich (St. Louis, MO) or Thermo Fisher Scientific (Waltham, MA).

In order to break any existing β-sheet structures and monomerize the protein, lyophilized Aβ was pretreated with hexafluoro-2-propanol at a concentration of 1 mg/ml for 40 mins until Aβ was fully dissolved. Aβ aliquots were transferred into glass scintillation vials, and hexafluoro-2-propanol was evaporated under vacuum overnight. Aliquots of dried peptide film were stored at −20°C. For an experiment, an Aβ aliquot was dissolved in freshly-made and filtered 60 mM NaOH and allowed to dissolve for 2 mins at room temperature. Tissue culture grade water was then added, and the vial was sonicated for 5 mins. The Aβ solution was then filtered with a 0.2-μm pore, 4-mm diameter syringe filter. Sterile phosphate buffered saline (PBS) was then added to the Aβ monomer solution yielding a final concentration of 222 μM with the NaOH:water:PBS ratio of 2:7:1. The Aβ solution was used immediately after preparation. HiLyte Aβ and FAM Scr Aβ were prepared in the same NaOH:water:PBS ratio solution to a stock Aβ concentration of 10 μM. All FCS experiments used 250 nM fluorophore-labeled Aβ.

### 2.2. Hydrogel Preparation

Rat tail collagen type I hydrogels were prepared to final concentrations ranging from 0.5 to 2 mg/ml. Cold 5x Dulbecco’s Modified Eagle’s Medium (DMEM) without phenol red, 7.5% sodium bicarbonate, sterile deionized water, and collagen were combined with PC12 cells to generate 3D substrates in black walled clear bottom well plates. The hydrogel solution (containing cells) was placed in a culture incubator for 20 mins to allow for gelation and then culture medium was added.

### 2.3. Cell Culture

PC12 cells (ATCC, Manassas, VA) (CRL-1721TM) were cultured on in collagen-coated flasks. Growth medium consisted of DMEM/F12 with L-glutamine and without phenol red, supplemented with 10% inactivated horse serum, 5% fetal bovine serum, and 20 μg/ml gentamicin. Experimental medium consisted of Neurobasal medium without phenol red, supplemented with 1% B27 and 10 μg/ml gentamicin. Phenol red and serum were avoided in the experiments because they are inhibitors of Aβ aggregation [65, 66].

### 2.4. Live/Dead Assay

PC12 cells were collected by trypsin treatment, and viability was determined by trypan blue staining. To remove serum, cells were resuspended in experimental medium, pelleted then resuspended again in experimental media. In a black-walled clear-bottom tissue culture treated 96-well plate, wells for 2D culture were collagen coated, and then PC12 cells were seeded at 15 × 10^3^ cell/cm^2^. For the 3D hydrogels, PC12 cells were mixed in collagen gel solution at a concentration of 500 cell/μl; the solution was then pipetted (30 μl) into the well plate and allowed to solidify. All wells were incubated in 200 µl warmed medium. The medium was not changed during the 72 hr experiment.

To determine cell viability, the Live/Dead mammalian cell kit (Invitrogen, Carlsbad, CA) was applied at a concentration of 4 μM Calcein AM (green-fluorescing live cell reporter) and 9 μM Ethidium homodimer-1 (EthD) (red-fluorescing dead cell reporter) and incubated at 37°C for 30 minutes. Images were taken on an IX81 Olympus inverted fluorescent microscope. A minimum of 100 cells were counted per well, two images per well, three wells per condition. The data is presented as percent viability, averaged between the three replicate experiments.

### 2.5. Thioflavin T

A black-walled clear-bottom 384 well plate (Costar) was sterilized under UV light for 15 minutes in a laminar flow hood. UltraPure grade Thioflavin T (ThT) (AnaSpec, Fremont, CA) was dissolved in deionized water at a concentration of 1 mM then filter-sterilized. Wells for 2D and 3D samples were prepared as above, but containing 20 µM ThT. The wells were sealed with black TopSeal-A membranes to prevent evaporation. The ThT experiment was analyzed on a Spectra Max M5 (Molecular Devices, San Jose, CA) spectrophotometer set to ex. 450 nm, em. 480 nm, at 37°C, taking reads every 30 mins for 72 hrs, reading from the bottom of the plate. Replicates were averaged, Aβ data was corrected with ThT control data, and corrected curves were normalized. Due to the stochastic nature of aggregation, representative curves are presented here.

### 2.6. Transmission Electron Microscopy

Samples for transmission electron microscopy (TEM) imaging were prepared as follows. Reconstituted 222 μM Aβ was diluted to 20 μM in experimental media. Timepoints were taken at 0, 24, 48, and 72 hrs. Samples were prepared on copper-supported carbon formvar grids (FCF200-Cu-TB) (EMS, Hatfield, PA) and stained with 0.2% uranyl acetate. Grids were imaged at 180-220 kx on an FEI Morgagni M268 100 kV Transmission Electron Microscope equipped with a Gatan Orius CCD camera.

### 2.7. Fluorescence Correlation Spectroscopy

#### 2.7.1. Theory

Fluorescence Correlation Spectroscopy (FCS) measures the fluctuations of fluorescence in a small, optically-defined confocal volume (~10^−15^ liter). These fluctuations are typically attributed to the fluorescent particles moving in and out of the volume with a statistical average residence time, τ_D_. The residence time is proportional to the hydrodynamic radius (R_H_) of the molecule. The fluctuations of detected photons inform the autocorrelation, G(τ), function defined as

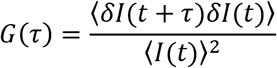

Where *δI*(*t*) = *I*(*t*) − 〈*I*(*t*)〉 is the fluorescence fluctuation determined from the measured fluorescence intensity, *I*(*t*), at time t, and the average intensity, 〈*I*(*t*)〉, over the period of measurement. The excitation laser, which is focused, is assumed to have a 3D Gaussian profile, with a characteristic radial dimension (w_0_) and a characteristic axial dimension (z_0_). For a solution of n noninteracting, freely diffusing fluorescent species G(τ) is given by:

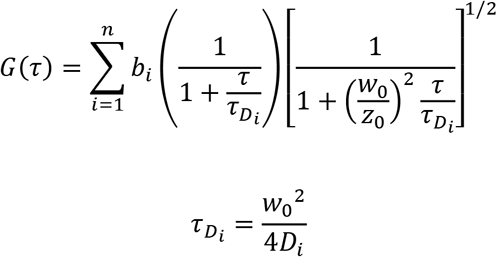

Here the D_i_ values are the n different values of diffusion constants and b_i_ are the relative fractions in brightness of these species. In practice, the radial and axial dimensions were determined using Alexa 488 dye in water where the diffusion coefficient (430 μm^2^/s) is known and was used to estimate the excitation volume for a 3D Gaussian beam [67].

#### 2.7.2. Methods

Neurobasal medium was used in preparing solution samples and contained 20 μM Aβ and 250 nM HiLyte Aβ. Collagen hydrogels were prepared as described with 20 μM Aβ and 250 nM HiLyte Aβ then pipetted into 0.8-mm deep hybridization chambers (PerkinElmer, Waltham, MA) on a borosilicate cover glass. Control samples were tested with 20 μM Scr Aβ and 250 nM FAM Scr Aβ.

The FCS measurements were performed using an Alba-FFS microscope-based system from ISS Inc. (Champagne, IL). The system is composed of: an Olympus IX81 inverted microscope equipped with a 60X/1.35NA oil immersion objective lens, a Prior Pro stage, three different lasers (450 nm, 488 nm, and 532 nm), two Hamamatsu photon multiplier tubes (PMTs) for photodetection, and two sets of computer-controlled scanned mirrors for imaging. In these measurements, only the 488-nm diode laser was used for excitation of the fluorophores Alexa 488 or fluorescently-labeled Aβ, and the emitted fluorescence was collected through confocal detection with a pinhole (< 50 mm) located in the image plane of the excited focused beam inside the sample. The emitted fluorescent beam was optically filtered further with (525/50nm) filter and then sent to a 50/50 beam splitter for detection by two PMTs positioned in a 90-degree angle configuration. The photocounts of both PMTs were continuously acquired and then computationally cross-correlated in order to eliminate the afterpulsing effect of a single PMT, which is typically noticeable at short delay times (< 10 ms).

Using Vista Vision software, two runs were carried out back to back collecting for 3 minutes each to generate the correlation function G(τ) for each sample at a time point. The two correlation functions were averaged, and the Scr Aβ correlation function was fit using the one-component model to determine the diffusivity of the monomer. Further, the measured time-correlation functions for Aβ were fit using the 2-component model where the size of species 1 was held constant at monomer diffusivity in order to derive the average aggregate diffusivity population of the second species. Additional refinement for fitting the correlation functions were also performed with the Maximum Entropy Method FCS (MEMFCS) thanks to a code gifted by Dr. S. Maiti (Tata Institute of Fundamental Research), allowing us to obtain the heterogeneous distribution of aggregate diffusivities at each time point [68].

Small molecules have a short delay time because they diffuse quickly through the volume, whereas large molecules have a long delay time because of their relatively slow diffusion through the volume. The 2-component model is intended to model two distinct molecular species in solution. For our samples, we held the monomer diffusivity constant as species 1 where the average diffusivity of aggregated species was identified by solving for species 2.

Fluorophore labeling of Aβ monomers inhibits aggregation due to the bulky groups sterically preventing proper monomer to monomer stacking [69]. Therefore, we used a ratio of 1:80 HiLyte 488-labeled Aβ to unlabeled Aβ, and FAM-labeled Scr Aβ to unlabeled Scr Aβ, to allow unhindered β-sheet stacking. Nanomolar fluorophore concentrations are also preferable in FCS in order for the detectors to monitor few individual fluorescent molecules in the confocal volume, enhancing hence the signal-to-noise of the fluctuations.

### 2.8. Statistical analysis

Data were analyzed for statistical significance with Prism v8 software (GraphPad). Cell viability data were analyzed with a general ANOVA with a post Tukey pairwise test which determined significant deviation from the population mean with a p-value <0.05 with 95% confidence. Error bars are the percent error of the mean population. The raw FCS experimental G(τ) curves, the 2-component model calculated G(τ) curves, and the MEMFCS calculated G(τ) curves were analyzed for significance using the 2-sample Kolmogorov-Smirnov test with 95% confidence.

## 3. Results

### 3.1. Toxicity of Aβ in 2D and 3D cultures

We initially hypothesized that 3D hydrogels would be better models in which to test Aβ toxicity because neurons have a more *in vivo* like phenotype in 3D compared to 2D cultures [23, 70]. As such, we expected Aβ to be more toxic to cells in 3D than in 2D. To begin to test this hypothesis, we examined the toxicity of PC12 cells when dosed with Aβ (pretreated to remove β-sheet structure) in both 2D and 3D collagen over a 72 hr period.

We acknowledge that the percent viability decreases for all samples over time, but it is important to note that the medium was not changed in order to better mimic the evolving population of Aβ species measured in the ThT and FCS experiments. Over 72 hrs, it is likely that cell waste accumulates and nutrients are depleted, thus explaining the decrease in cell viability in all conditions. The number of viable cells cultured in 2D with 20 μM pretreated Aβ decreased significantly by 24 hrs where the cell viability was 49% (p-value 0.004); at 48 hrs and 72 hrs the cell viability was only 16% (p-value <0.001) and 12% (p-value <0.001) respectively (Figure 1a). When cells were encapsulated within 3D collagen hydrogels, treatment with Aβ did not affect cell viability (Figure 1b). Representative fluorescence microscopy images of stained cells at 0 hrs, 24 hrs, and 48 hrs in 2D and 3D culture are shown in Figures 1c and 1d respectively. In the presence of Aβ, the cell death (red staining) at 48 hours in 2D culture is striking (Figure 1c), while no notable increase in cell death is observed in the Aβ-treated 3D cultures (Figure 1d).

**Figure 1.**
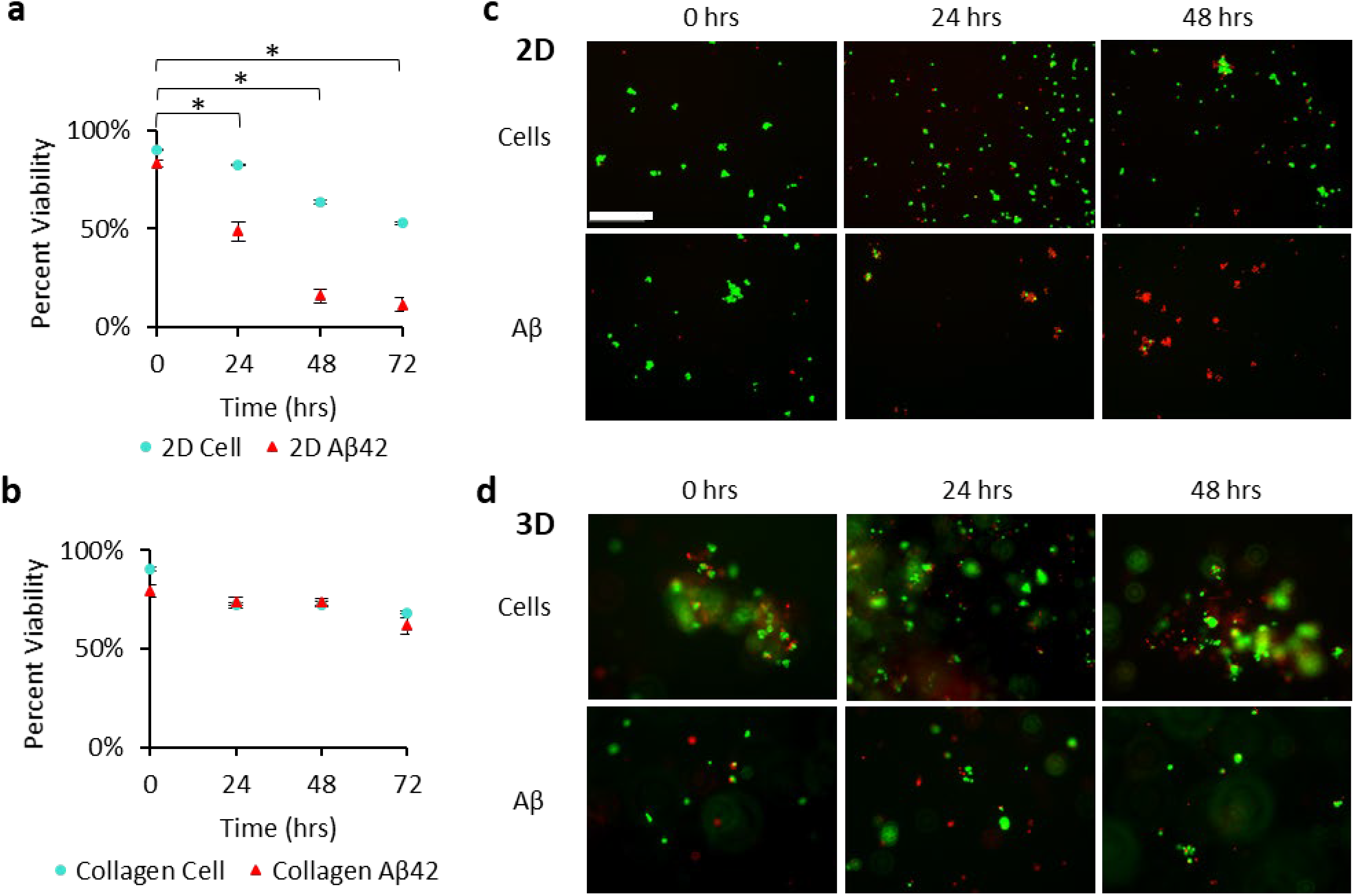
PC12 cell percent viability in 2D and 3D culture. Using a Live/Dead assay without Aβ (**a** & **b**; turquoise, •), or with 20 μM Aβ (**a** & **b**; red, ▴). Cells were cultured on 2D collagen I coating (**a**), or encapsulated within a 1 mg/ml collagen hydrogel (**b**). Cell were imaged in 2D (**c**) and 3D (**d**), without Aβ and with Aβ, where live cells appear green (Calcein-AM) and dead cells appear red (EthD). Significant differences were seen in (**a**) 2D culture in the presence of Aβ at 24, 48, and 72 hrs signified by (*). Statistics used n = 4. P values at significantly different times in 2D culture: 24 hrs (0.004), 48 hrs (<0.001), and 72 hrs (<0.001). Error bars are the percent error of the mean population. Scale bar is 200 μm; all micrographs are the same magnification.

### 3.2. TEM imaging of Aβ in solution

Given the well-known correlation between Aβ structure or aggregation state and toxicity, we sought to confirm that the Aβ used in these studies aggregated as expected and that the surprising cell viability results were not the result of some anomalous aggregation process. To this end, we performed TEM imaging of the pretreated Aβ at time points after dissolution relevant to the viability studies.

Representative TEM micrographs are shown in Figure 2. At time zero, unstructured protein globules of various sizes were identified (Figure 2a). The smallest structures have a diameter of ~3.6 nm, which correlates well to the size of Aβ monomer (hydrodynamic radius of 1.8 nm) [71]. Larger globular structures are 6 – 50 nm in diameter and lack any fibrillar structures. At 24 hrs, thin filamentous aggregates (~3 nm diameter) are seen to extend from the unstructured globules (Figure 2b). By 72 hrs, thick Aβ fibrils are present and associate with each other into structures ~13 nm in diameter (Figure 2c).

**Figure 2.**
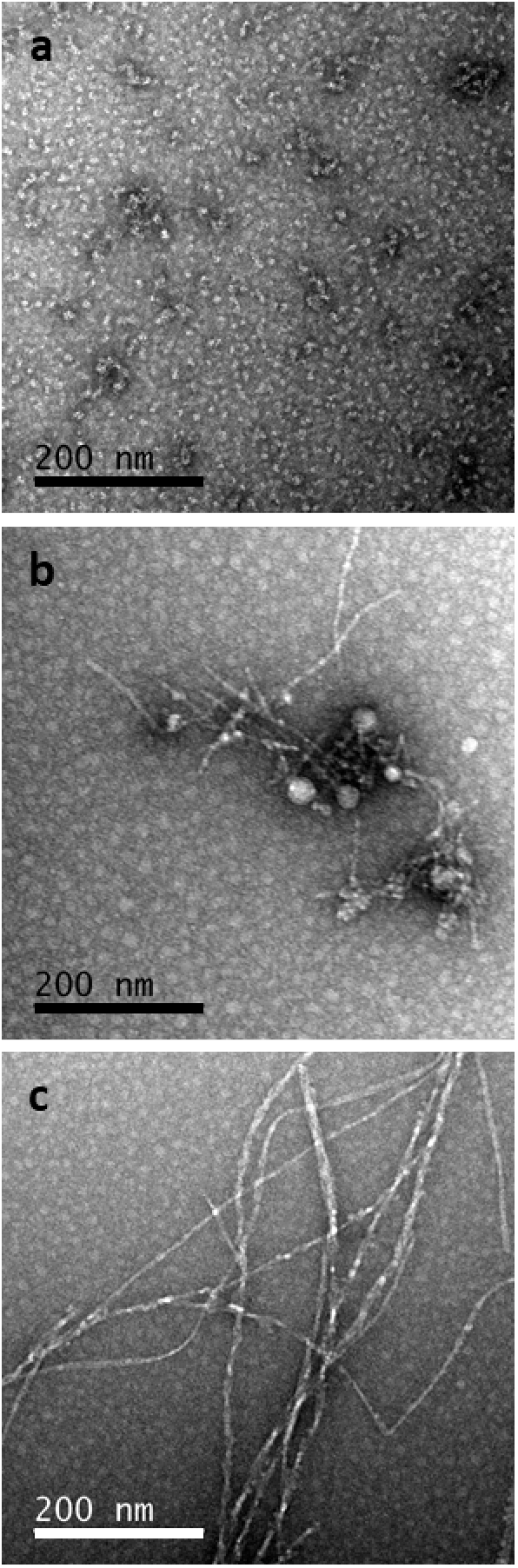
TEM images of Aβ in solution. Pretreated Aβ (20 μM) was stained with 0.2% uranyl acetate at time 0 hrs (**a**), 24 hrs (**b**), and 72 hrs (**c**). Scale bar is 200 nm.

### 3.3. Aβ aggregate diffusivities by FCS

Given the inherent challenges of imaging hydrogels using TEM, we utilized FCS to infer relative Aβ aggregate size from the diffusivity of fluorescently-labeled Aβ species. Diffusivity scales inversely to the radius of a particle. Therefore small diffusivity values correspond to large particles.

Non-aggregating Scr Aβ was measured in solution and hydrogels as a monomer control. The diffusivity of these Scr Aβ monomers in solution was determined to be 175 μm^2^/s, whereas the diffusivity in the hydrogel was 129 μm^2^/s (Figure 3a & b). As points for comparison, the diffusivity of Aβ monomer has been reported to be 180 μm^2^/s in solution and 62.3 μm^2^/s in brain tissue [72]. In solution, the diffusivity of the average Aβ aggregate population (determined using the 2-component model) is ~9x slower than the monomer for up to 6 hrs (Figure 3a). In hydrogels, the diffusivity of the average Aβ aggregate population is ~150x slower than a monomer for up to 4 hrs (Figure 3b). The 2-component calculated G(τ) of solution is significantly different than the calculated G(τ) of the collagen gel with a p-value of 0.0003. Additionally, the raw experimental G(τ) of solution is significantly different from the raw experimental G(τ) of the collagen gel with a p-value of <0.0001.

**Figure 3.**
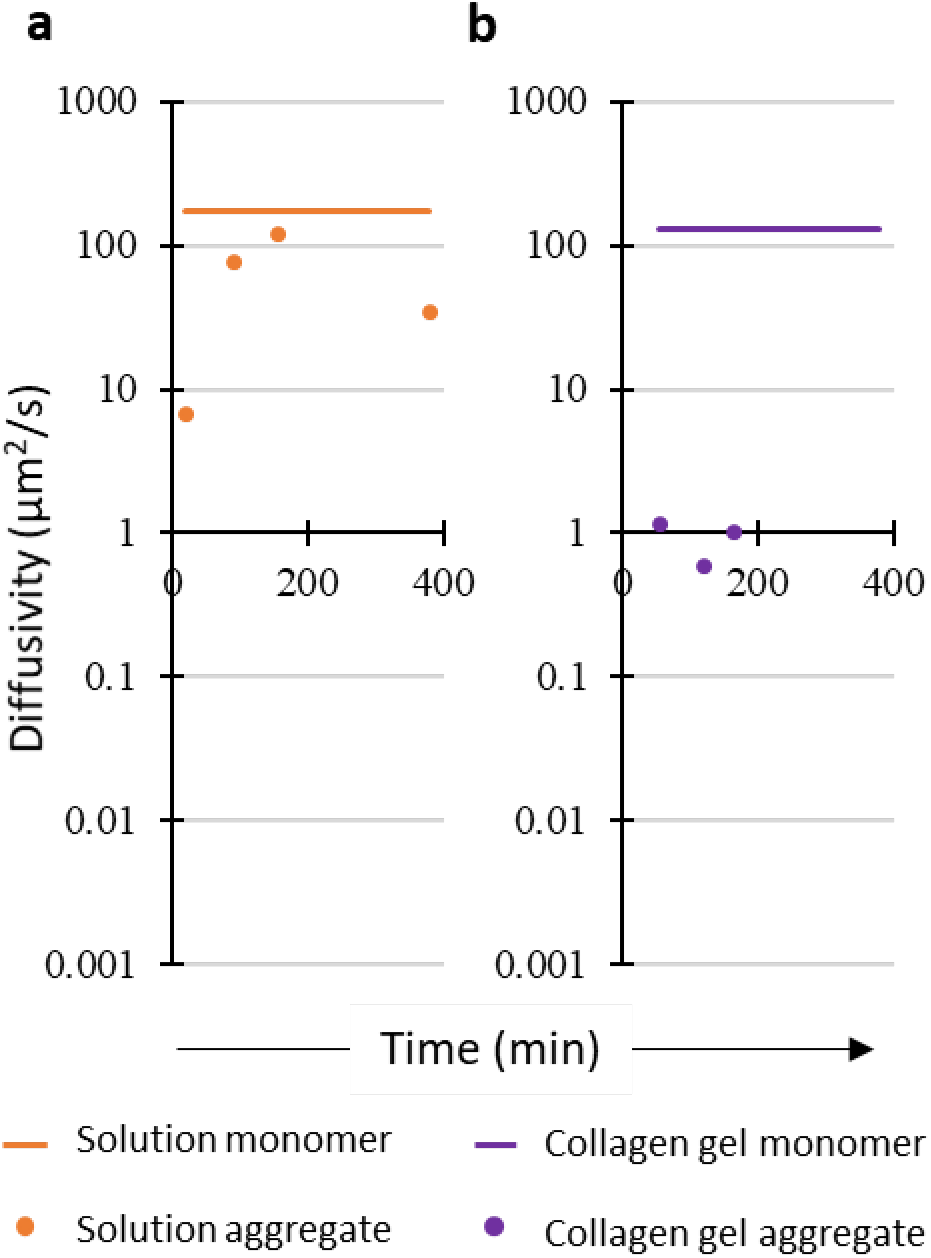
FCS correlation functions of Aβ were fit with a 2-component model. Samples of 20 μM Aβ with 250 nM HiLyte Aβ in solution (**a**; orange, •) and in collagen hydrogel (**b**; purple, •) were analyzed. Species 1 was held constant at the diffusivity of the Scr Aβ control assumed to be monomer (solid line). Species 2 was calculated and represents the average Aβ aggregate diffusivity population (•). There is a significant difference between the 2-component calculated G(τ) of solution and collagen gel, p-value: 0.0003 (solution n=8; collagen gel n=7).

The correlation functions were also analyzed using the MEMFCS program, which attempts to determine the distribution of size of the aggregating solutions. A distribution of multiple diffusivity populations of Aβ aggregates and their relative fractions were modeled. In solution, Aβ diffusivity values have a single broad distribution with an average peak diffusivity of 90 μm^2^/s (Figure 4, solution). The peak diffusivity of Aβ in solution is ~2x slower than the Scr Aβ diffusivity, suggesting an Aβ population predominately composed of dimers. In the hydrogel, Aβ has a peak diffusivity of 60 μm^2^/s, which is ~2x slower than the Scr monomer (129 μm^2^/s). However, in contrast to the solution samples that only have one diffusivity peak, the hydrogel sample data show a small secondary diffusivity peak as early as 5 mins after addition of Aβ to the hydrogel and persists throughout the measurement period (up to 4 hrs) with diffusivity values in the range of 0.2 μm^2^/s to 3 μm^2^/s, or between 650x to 45x slower than Scr Aβ (Figure 4, collagen). The MEMFCS calculated G(τ) of solution is significantly different from the collagen gel calculated G(τ) with a p-value of 0.0004.

**Figure 4.**
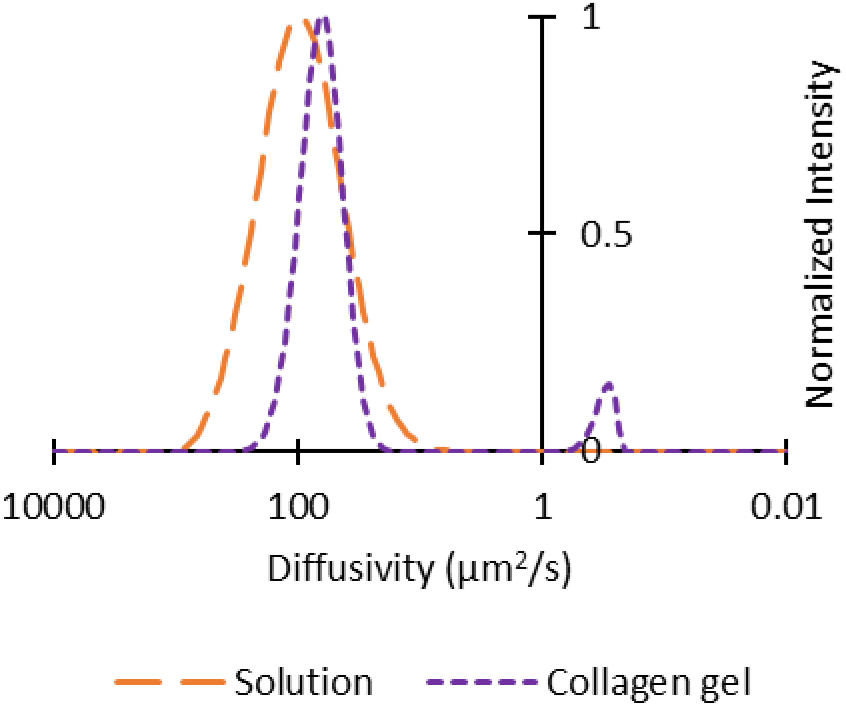
FCS correlation function was fit using the MEMFCS program gifted by Dr. S. Maiti [68]. Solution timepoints were collected over 8 hrs; a representative curve is shown (long dash, orange). Collagen hydrogel timepoints were collected over 4 hrs; a representative curve is shown (short dash, purple). There is a significant difference between the MEMFCS calculated G(τ) of solution and collagen gel, p-value: 0.0004 (solution n=6; collagen n=6).

Both analysis methods of the FCS data indicate that Aβ aggregates differently in 2D compared to 3D. Based on these data, a rough estimate of aggregate species size in 3D is ~25x to 200x larger in diameter than the Aβ species detected in 2D.

### 3.4. ThT fluorescence as a measure of Aβ aggregation kinetics

While FCS is a powerful technique to measure diffusivities of molecules in complex media, it is difficult to get direct information on the structure or morphology of an aggregated species. Therefore, we used a ThT fluorescence assay as an additional measure of Aβ aggregation.

Representative curves of ThT fluorescence vs. time are shown in Figure 5 for Aβ aggregation in solution and collagen hydrogels. In solution, fibrillar Aβ aggregation (signified by ThT fluorescence) has a lag phase during the first ~20 hrs, followed by rapid aggregation (Figure 5, orange). In hydrogels, however, fibrillar aggregation does not show a lag phase, and instead, fluorescence steadily increases from the initiation of the experiment (Figure 5, purple). Depending upon the supplier and lot of Aβ tested, these curves lag time, and fluorescence intensity varied, but the qualitative differences between fibril Aβ aggregation in solution and the collagen hydrogel persisted; fibril aggregation was accelerated in the hydrogel compared to the solution.

**Figure 5.**
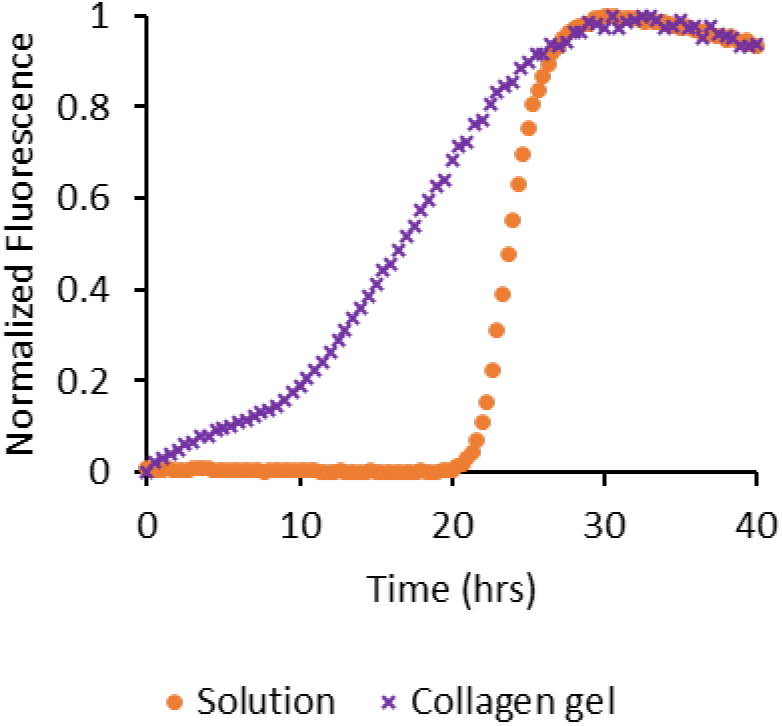
ThT fluoresces only when bound to stacked β-sheet structures. Shown are representative data of ThT fluorescence due to Aβ aggregation in solution (orange; •) and collagen hydrogel (purple; 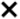).

In solution samples, the duration of the aggregation lag phase varied with Aβ concentration: greater Aβ concentrations correlated with shorter lag phases of fibril aggregation in solution. The lag phase lasted 7.5 hrs for 10 μM Aβ, 4 hrs for 25 μM Aβ, and only 1 hr for 50 μM Aβ (Figure 6a). This effect was also noted for hydrogels, but on a much faster timescale: the lag phase was 1 hr for 10 μM Aβ, and nonexistent for 25 μM and 50 μM Aβ (Figure 6b). The rate of fibril aggregation was also influenced by the collagen concentration of the hydrogel: at collagen concentrations of 0.5, 1, 1.5 and 2 mg/ml, the lag phases were observed to be 13.5 hrs, 9.5 hrs, 9 hrs, and 6.5 hrs, respectively (Figure 6c).

**Figure 6.**
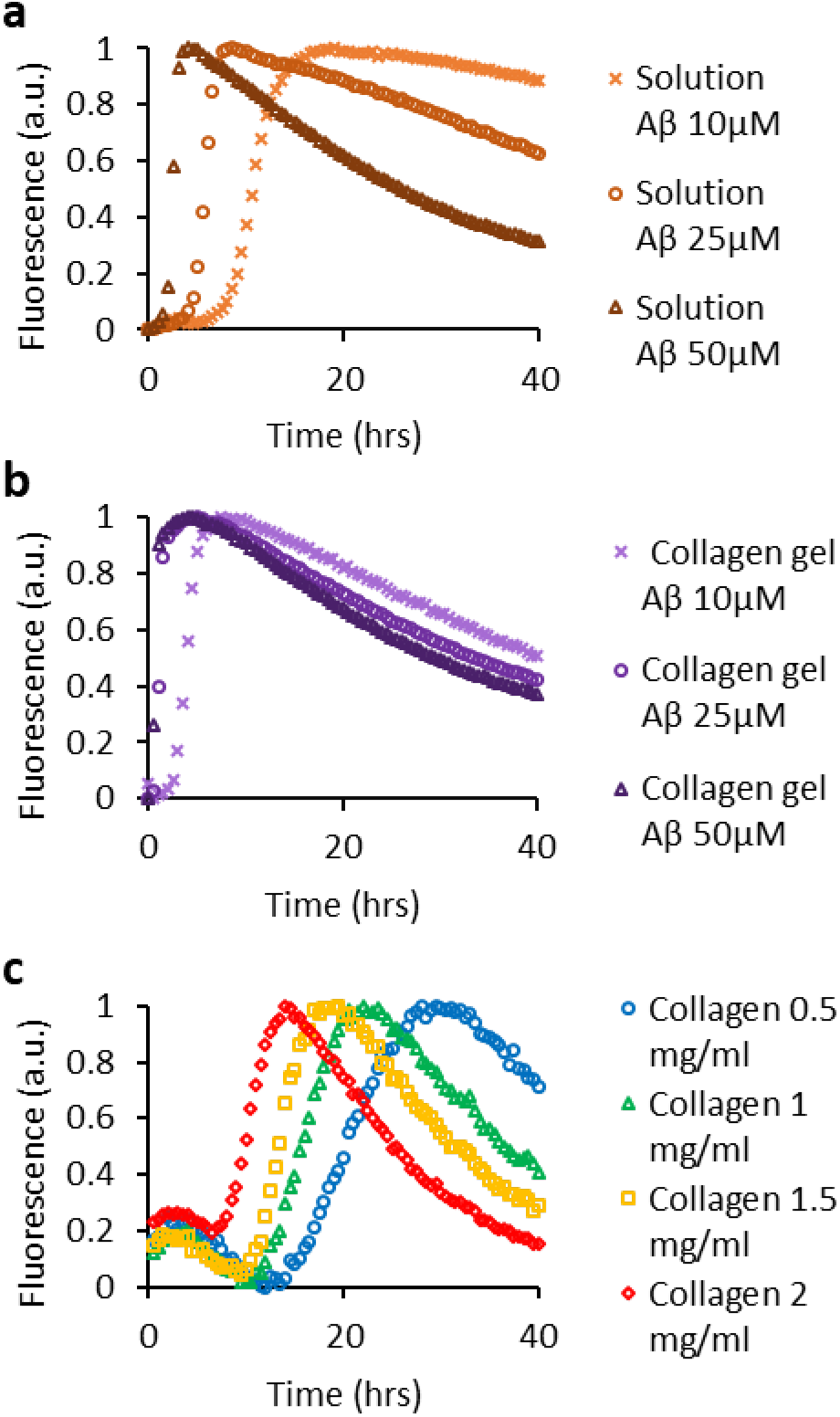
Aβ and collagen concentration impact ThT aggregation kinetics. ThT aggregation kinetics comparing Aβ concentrations (10 μM, light 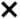; 25 μM, medium 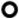; 50 μM, dark 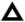) in solution (**a**, orange) and collagen hydrogel (**b**, purple). In **c**, Aβ aggregation kinetics is compared in hydrogels with collagen concentrations of 0.5 mg/ml (blue; 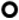), 1 mg/ml (green; 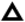), 1.5 mg/ml (yellow; 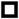), and 2 mg/ml (red; 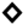).

## 4. Discussion

For almost three decades, researchers have recognized a link between Aβ and AD and pursued drug candidates capable of preventing Aβ aggregation as new therapeutic strategies for AD [73–75]. These interactions were measured in cell culture experiments but were associated with little or no efficacy in pre-clinical testing and clinical trials [7, 9, 76–79].

There is a growing appreciation that cells grown *in vitro*, on rigid polystyrene tissue culture surfaces, behave very differently than cells *in vivo*, and that the stiffness of the surface upon which cells are grown can have profound epigenetic effects on cells that influence their phenotype [80, 81]. In addition, many investigators have observed that morphologies of neuronal cells are strikingly more similar to those expressed *in vivo* in 3D vs. 2D culture [22, 23, 82, 83]. Thus, the use of 3D cell culture models for AD drug development may lead to a more positive correlation between successes *in vitro* and successes in clinical outcomes.

The preponderance of literature reports indicates that Aβ is toxic to cells in culture only when aggregated and that it is the oligomeric or intermediate-sized aggregation species that are the most toxic species, with large fibril aggregates being less toxic [57, 62]. These findings were reported for cells in traditional culture on flat rigid substrates such as polystyrene or glass. To our knowledge, no one has explored Aβ aggregation kinetics and cellular toxicity in a 3D cell culture model.

In the work presented here, our 2D cell culture of PC12 cells demonstrated the expected cytotoxic nature of Aβ aggregates. However, when PC12 cells and Aβ were encapsulated within 3D collagen hydrogels, the cytotoxic effects were attenuated. We supposed that this effect could be due to one of two effects, either (1) the 3D environment altered the phenotype of the PC12 cells, making them less susceptible to Aβ, or (2) the 3D environment altered the aggregation of Aβ. Given that PC12 cells and other neuron-like cells take on a phenotype more closely associated with an *in vivo* neuron in 3D [22, 23, 84], we assumed that our first hypothesis was less probable. Therefore, in this work, we set out to investigate how the 3D collagen hydrogel environment affects Aβ structure and aggregation.

We used a combination of biophysical methods to investigate Aβ structure and aggregation both in solution and in the 3D collagen hydrogel. TEM micrographs of pretreated Aβ that underwent aggregation in solution confirm that the structures seen during our experiments are typical of those seen by others [85, 86]. The pretreatment process removed any observable fibril or protofibril structures (Figure 2a). FCS and ThT measurements at these early time points (<24 hrs) suggest that only small, rapidly diffusing species remained after Aβ pretreatment; the samples were devoid of any extended β-sheet structures (Figures 3a, 4, and 5, respectively). These results are consistent with literature reports for Aβ pretreatment via this method [59, 71]. Shortly after the dissolution of Aβ, the Aβ in solution appeared to form dimer or similar diffusing species (Figure 4, solution), and other somewhat larger aggregated species with diffusivity values on the order of 20 μm^2^/s (Figure 3a). The observance of monomers and dimers in equilibrium at early stages of aggregation of Aβ in solution has been seen via other methods including chromatography [87, 88] and hydrogen exchange mass spectrometry and others [89, 90]. The observance of larger aggregated species in Aβ solutions that are non-fibril (i.e., they do not bind ThT; Figure 5), such as the more slowly diffusing species seen in FCS (Figure 3a) and TEM micrographs (Figure 2b), is consistent with the formation of the toxic oligomer species. While there is considerable debate concerning the actual size and structure of the most toxic species of aggregated Aβ, most consider that this species is smaller than a fibril [91], and by some accounts, the diffusivity of the toxic Aβ species is on the order of 20 μm^2^/s [63, 92]. Aβ aggregated for 72 hours shows typical fibril structures via TEM (Figure 1c), that bind ThT (Figure 5), as would be expected for large extended β-sheet structures [93]. Given the typical aggregation pathway, we observed of Aβ in solution – one that included detectable oligomers that are suggested to be the most toxic species – the viability results observed in 2D cell culture (Figure 1a & 1c) were expected.

The same careful biophysical examination of Aβ aggregation in 3D collagen hydrogels demonstrates that aggregation does not proceed through the same pathway as in solutions. Even at early times during aggregation, no slowly diffusing species (diffusivity ~20 µm^2^/s) were observed (Figures 3b, 4). At these early times during aggregation, only a small unaggregated species and a very large/slow diffusing species were observed; the larger species diffused 10x to 100x more slowly than the oligomer observed in solution (Figures 3, 4). Given the ThT fluorescence seen even at these early times (Figure 5), this larger species may be a fibril form of Aβ. We hypothesize that we did not observe any notable Aβ toxicity in 3D culture because Aβ aggregation proceeded through a pathway that did not include the formation of any detectable oligomeric species that are typically associated with Aβ toxicity.

While we are not aware of any other studies of Aβ aggregation in a 3D environment, the effect of crowding and confinement on protein folding and aggregation has been previously studied [41, 48]. Biomaterial scaffolds are composed of self-associating or crosslinked structures that exert an excluded volume effect on soluble molecules that are entrapped within. The excluded volume effect increases the local concentration of soluble molecules, increasing the chance of protein-protein interactions. The reduced volume also confines the available space a protein may use to fold, promoting a more compact, low entropy conformation. Uncrosslinked crowding molecules such as PEG, Ficoll, and bovine serum albumin stabilize protein folding into functional states or accelerate aggregation in aggregation-prone proteins (e.g., lysozyme, α-synuclein, and β2-microglobulin) [44, 45, 48]. Confinement has a greater stabilization effect than crowding but is seldom used to study the biophysics of proteins [39, 94, 95]. We suggest that in our studies, the confinement provided by the 3D collagen hydrogel resulted in increased Aβ-Aβ interactions, thus leading to the observed faster aggregation kinetics and the lack of slower diffusing species. If this were the case, then we would expect to see that higher concentrations of Aβ shortened the lag time to the onset of rapid aggregation due to the greater probability of nucleation of aggregation. This is indeed what we observed (Figure 6a and b). At every concentration tested, the lag phase until the onset of rapid aggregation was significantly shorter in 3D environments and shorter at higher Aβ concentrations. The acceleration of aggregation of Aβ with increasing concentrations of collagen in the 3D hydrogels (Figure 6c) is consistent with the hypothesis as well; higher collagen concentrations would lead to smaller void spaces in the 3D collagen hydrogel, thus greater confinement, increased Aβ-Aβ interactions, and acceleration of Aβ aggregation.

When Aβ aggregation is nucleated in a confined environment, the β-sheet structures that form are more stable vs. aggregation as free molecules in solution, resulting in faster filament growth. Such stabilization of β-sheet structured Aβ aggregates that occurs in collagen hydrogels should result in a shift in the chemical equilibrium of aggregate species. In solution, the equilibrium composition of Aβ species is expected to be a distribution of unstructured monomer-dimer – multimer species. In confined collagen hydrogels, however, we hypothesize (and our data support) that the equilibrium is shifted to stable β-sheet aggregates and away from the relatively unstable/unstructured presumed toxic oligomeric Aβ species.

We propose that the attenuation of Aβ toxicity observed in the 3D environment is due to the shift in aggregation kinetics of Aβ away from the smaller toxic oligomer. However, other explanations are also plausible. Aggregated Aβ species may have hindered diffusion in the 3D collagen hydrogel. The FCS data suggest that at least at early times the Aβ species do diffuse within the hydrogel. In addition, the mesh size of the 3D collagen hydrogels used in this study was ~10 μm, one to two orders of magnitude larger than reports of sizes of Aβ toxic oligomers [96, 97]. Therefore, we do not conclude that hindered diffusion of Aβ within the hydrogel contributed significantly to the attenuation in Aβ toxicity observed in 3D compared to 2D environments. It is also possible that the 3D environment resulted in some unexpected phenotypic change in the PC12 cells that we cannot rule out in the studies performed in this work.

The shift in the equilibrium of Aβ species away from smaller potentially toxic species to larger extended β-sheet species in confined environments observed in our studies may suggest similar phenomena *in vivo* tissues. At the very least, our studies suggest that Aβ aggregation kinetics are fundamentally different in 3D structures as compared to in the solution phase of 2D culture, making a direct translation of results from a 2D culture model to a more *in vivo*-like 3D culture model difficult. However, this relatively simple collagen hydrogel cannot recapitulate the complexity of the *in vivo* environment. We can only speculate that the high percentage of failed Aβ targeting drugs in clinical trials may be due to the drug discovery models missing essential properties such as confinement and dimensionality. Knowing the biophysical impact of confinement stabilizing toxic Aβ aggregate species, advanced 3D *in vitro* models may be developed to investigate AD pathology and be implemented in AD drug development.

## Acknowledgments

The authors would like to thank Dr. Tagide deCarvalho for her assistance with TEM imaging and Dr. S. Maiti (Tata Institute of Fundamental Research) for sharing the Maximum Entropy Method program for FCS.

## Funding

This work was supported by funding from the National Science Foundation (NSF) [EAGER CBET-1447057] and the National Institute of Health (NIH) [R01GM117159]. NSF provided support for TAG to contribute to this project through their Independent Research and Development program. Any opinion, findings, and conclusions or recommendations expressed in this material are those of the author(s) and do not necessarily reflect the views of the National Science Foundation.

